# Aperiodic measures of neural excitability are associated with anticorrelated hemodynamic networks at rest: a combined EEG-fMRI study

**DOI:** 10.1101/2021.01.30.427861

**Authors:** Michael S. Jacob, Brian J. Roach, Kaia Sargent, Daniel H. Mathalon, Judith M. Ford

## Abstract

The hallmark of resting EEG spectra are distinct rhythms emerging from a broadband, aperiodic background. This aperiodic neural signature accounts for most of total EEG power, although its significance and relation to functional neuroanatomy remains obscure. We hypothesized that aperiodic EEG reflects a significant metabolic expenditure and therefore might be associated with the default mode network while at rest. During eyes-open, resting-state recordings of simultaneous EEG-fMRI, we find that aperiodic and periodic components of EEG power are only minimally associated with activity in the default mode network. However, a whole-brain analysis identifies increases in aperiodic power correlated with hemodynamic activity in an auditory-salience-cerebellar network, and decreases in aperiodic power are correlated with hemodynamic activity in prefrontal regions. Desynchronization in residual alpha and beta power is associated with visual and sensorimotor hemodynamic activity, respectively. These findings suggest that resting-state EEG signals acquired in an fMRI scanner reflect a balance of top-down and bottom-up stimulus processing, even in the absence of an explicit task.

**HIGHLIGHTS:**

- Periodic and aperiodic EEG parameters associated with distinct resting-state networks
- Increases in aperiodic power associated with an auditory-salience-cerebellar network
- Decreases in aperiodic power associated with prefrontal regions
- Global neural excitability may reflect stimulus processing or arousal attributable to the uniqueness of the resting-state MR-scanner environment

## 1. INTRODUCTION

Despite the name, “resting-state” recordings are associated with widespread and pronounced neural and metabolic activity (Northoff et al., 2010; Raichle, 2015; Raichle et al., 2001). Metabolic and hemodynamic studies, primarily using PET and fMRI, have consistently identified a network of brain regions active in the absence of an explicit task, known as the default mode network (DMN, for review see: Buckner et al., 2008; Mak et al., 2017; Raichle, 2015). This activity has been termed “dark energy,” (Capolupo et al., 2013; Raichle, 2006; Zhang and Raichle, 2010) seemingly unaccounted for by external stimulus processing, and instead, linked to internal thought processes within the DMN. However, hemodynamic activity is an indirect measure of neural activity (Hillman, 2014; Logothetis and Wandell, 2004; Mishra et al., 2021) representing a mixture, or balance of incoming excitatory and inhibitory (E/I) dendritic currents (Xu, 2015). Ongoing, seemingly “spontaneous” fluctuations in neural excitability are critical for neurodevelopment (Blankenship and Feller, 2010) and persist into adulthood despite high metabolic demand (Zhou and Yu, 2018). Moreover, the relationship between ongoing electrophysiologic activity and resting-state networks (commonly identified from hemodynamic and metabolic signals, such as fMRI and PET) remains obscure (Foster et al., 2016).

Resting-state neural activity as measured by EEG and ongoing spontaneous activity at the level of cellular neurophysiology is both periodic and aperiodic (Buzsaki, 2006). The broadband, aperiodic component typically lacks a dominant temporal scale, a property often referred to as ‘scale-free’ (He, 2014; He et al., 2010) or ‘1/f-like’ (Donoghue et al., 2020). Scale-free phenomena are ubiquitous in biological systems (Eke et al., 2002; Gisiger, 2001; Werner, 2010) including the brain: from the fractal patterns present in dendritic trees (Ristanović et al., 2014) (spatially scale-free) to the spiking activity of neurons (Petermann et al., 2009) (temporally scale-free). Scale-free EEG may reflect criticality dynamics of neural “avalanches” (He, 2014; R. Chialvo, 2004; Johnson et al., 2019); that predict processing speed (Ouyang et al., 2019) and behavioral variability (Palva et al., 2013). Despite receiving the most attention in the EEG literature, oscillatory and periodic activity is not the primary basis for scalp EEG (Bullock et al., 1995), and reflects a much smaller proportion of the total EEG power than aperiodic activity (Figure 2A). Scale-free EEG has been hypothesized to be analogous to the DMN: a signature of task disengagement (Freeman et al., 2009), from which EEG rhythms emerge to support cognition (Freeman, 2006).

Aperiodic EEG activity can be characterized by estimating the slope and/or intercept of a linear fit to EEG spectra after log-log transformation of frequency and power. The aperiodic exponent,**β**, in the 1/*f*^**β**^ model, is equivalent to the negative slope in log-log space and is linked to neuronal activation (Podvalny et al., 2015), aging (Voytek et al., 2015) and development (He et al., 2019). The aperiodic exponent has been described as ‘electrophysiological marker of arousal level’ (Lendner et al. 2020) as it indexes a range of arousal states including sleep (Ma et al., 2018), depth of anesthesia (Colombo et al., 2019), and drug induced states (Muthukumaraswamy and Liley, 2018), consistent with the theory that aperiodic parameters index widespread brain activity including network excitability and E/I balance (Gao et al., 2017). The neurometabolic correlates of periodic EEG activity have been examined using simultaneous EEG-fMRI (Laufs, 2008). Much of this work has utilized one or two EEG electrodes (Laufs et al., 2003b), an average of all EEG electrodes (Mantini et al., 2007) or focused investigations of DMN and alpha power (Bowman et al., 2017; Goldman et al., 2002; Laufs et al., 2003a; Mayhew and Bagshaw, 2017; Moosmann et al., 2003; Scheeringa et al., 2012).

Aperiodic and related measures have also been applied to the BOLD signal (Ciuciu, 2012; Herman et al., 2011; Maxim et al., 2005; Nagy et al., 2017; Thurner et al., 2003), with some preliminary evidence suggesting that aperiodic EEG activity may underlie the fMRI global signal (Wen and Liu, 2016a). The scale-free organization of EEG is thought to be metabolically optimal (Roberts et al., 2014) and emergent from resource constraints (Burroni et al., 2017), highlighting the importance of considering scale-free neural activity in the context of metabolic analyses, such as that can be accomplished with fMRI.

Using simultaneous EEG-fMRI in healthy participants at rest with their eyes open, we investigated the possibility that aperiodic EEG and the hemodynamically defined DMN, two dominant signatures of brain activity, were associated with each other. We first extracted aperiodic and periodic components of the EEG spectra using a recently developed algorithm to parameterize neural power spectra (Donoghue et al., 2020). We then used these parameters as predictors in a whole-brain, parametric modulation analysis to identify hemodynamic activity correlated with aperiodic and periodic EEG activity. Contrary to our initial hypothesis, we found that aperiodic EEG activity shows minimal association with activity in the DMN but is instead positively correlated to an auditory-salience-cerebellar network and negatively correlated to prefrontal executive control networks. Residual periodic activity in alpha and beta bands is associated with reductions in visual and sensorimotor areas, with smaller contributions of gamma activity to auditory regions.

## 2. METHODS

### 2.1 Participants

Data are reported from 46 healthy participants (ages 18-69), without medical, neurologic or psychiatric illness. All participants were screened using the Structured Clinical Interview for DSM-IV (SCID-IV) (First et al., 1995) and recruited through advertising and word of mouth. Exclusion criteria included any current or past psychiatric disorder based on the SCID-IV, any significant medical or neurological illness, head injury resulting in loss of consciousness, substance abuse in the past 3 months or lifetime substance dependence (except nicotine/caffeine). Study procedures were approved by the University of California at San Francisco and the San Francisco Veterans Affairs Medical Center Institutional Review Boards (FWA #00000068). All participants provided written informed consent prior to the participation in the study. Demographic data are presented in Table 1.

**Table 1.** Group Demographic Data^a^

| **Table 1.** | **Participant Data** |
| --- | --- |
| **Number of participants** | 46 |
| **Age (years)** | 38.3 (15.1) [18 - 69] |
| **Gender** | 9W, 37M |
| **Average Parental SES<sup>b</sup>** | 34.6 (18.6) |
| **Handedness<sup>c</sup>** | 41R, 2L, 3A |
| **Estimated IQ<sup>d</sup>** | 111.3 |

<sup>a</sup>Note: Values are given as number of men (M) and women (W) gender, right (R), left (L), and ambidextrous (A) handedness, and antipsychotic type. Group means with the standard deviation for age, parental socioeconomic status (SES), intelligence quotient (IQ),<sup>b</sup> The Hollingshead (1975) four-factor index of parental socioeconomic status (SES) is based on a composite of maternal education, paternal education, maternal occupational status, and paternal occupational status. Lower scores represent higher SES.
<sup>c</sup> The Crovitz-Zener (1962) questionnaire was used to measure handedness and categorize it as right (R), left (L), or ambidextrous (A).R, L, or A.
<sup>d</sup> The Wechsler Adult Intelligence Scale (WAIS-III) full-scale intelligence quotient (FSIQ) was estimated based on the Wechsler Test of Adult Reading (WTAR) for native English-speaking subjects. Estimated IQ values are missing from 3 participants.

### 2.2. fMRI Recording Procedures

Simultaneous EEG-fMRI data were acquired at rest, with participants instructed to keep their eyes open and fixated on a white cross displayed on a black screen for 6 minutes. Structural and functional MRI data were collected using a 3T Siemens Skyra scanner. The details of our EEG-fMRI recording procedures have been previously described (Ford et al., 2016). The structural imaging protocol was a magnetization-prepared rapid gradient-echo (MPRAGE) T1-weighted high-resolution image (2300 ms TR, 2.98 ms TE, 1.20 mm slice thickness, 256 mm field of view, 1.0 × 1.0 × 1.2 voxel size, flip angle 9°, sagittal orientation, 9:14 min). The fMRI protocol was an AC-PC aligned echo planar imaging (EPI) sequence (2000ms TR, 30ms TE, flip angle 77°, 30 slices collected sequentially in ascending order, 3.4 × 3.4 × 4.0 mm voxel size, 182 frames, 6:08 min).

### 2.3 fMRI Preprocessing

We used Statistical Parametric Mapping 8 (SPM8; http://www.fil.ion.ucl.ac.uk/spm/software/spm8/) for image preprocessing. Motion correction was performed via affine registration and realigned to the first image of the run. Images were slice-time corrected with respect to the middle slice to adjust for timing differences of individual slices within each TR. We implemented aCompCor (anatomic component based noise correction, (Behzadi et al., 2007), a principal components-based approach to noise reduction of fMRI time-series data. Briefly, aCompCor derives principal components from the time series of voxels within “noise” regions of interest (ROIs) defined on from white matter and cerebrospinal fluid (CSF) parcels from participants’ high-resolution T1-weighted images. These data are subjected to PCA, to identify “noise” components comprising weighted averages of white matter and CSF voxel time series.

### 2.4 EEG Acquisition

Continuous EEG data were collected from 32 scalp sites (Fp1, Fp2, F3, F4, C3, C4, P3, P4, O1, O2, F7, F8, T7, T8, P7, P8, Fz, FCz, Cz, Pz, Oz, FC1, FC2, CP1, CP2, FC5, FC6, CP5, CP6, TP9, TP10, POz) with an additional electrode placed on the back to monitor ECG. We used sintered Ag/AgCl ring electrodes in an MR-compatible electrode cap from Brain Products (Munich, Germany) according to the 10-20 system of electrode placement. Electrode impedances were kept below 10 kΩ. The nonmagnetic, battery powered EEG amplifier was placed in the scanner bore behind the head coil and stabilized with sandbags. The subject’s head was immobilized using cushions. All 32 channels were recorded with FCz as reference and AFz as ground. The data were recorded with a bandpass filter of 0.1–250 Hz and digitized at a rate of 5 kHz with 0.5 μV resolution (16 bit dynamic range, 16.38 mV).

### 2.5 EEG Preprocessing

Our EEG preprocessing procedures utilized here have been previously described in detail (Ford et al., 2016). Data collected from an electrocardiogram (ECG) electrode were used to identify heartbeat artifacts in the continuous EEG signal using a semi-automatic heartbeat detection routine in Brain Vision Analyzer. MR gradient and cardioballistic artifacts were subsequently removed using artifact subtraction (Allen et al., 2000) implemented in Brain Vision Analyzer 2.0 software (BrainProducts) and then down-sampled to 250 Hz. Canonical correlation analysis (CCA) was used to remove broadband or electromyographic noise from single trial data (De Clercq et al., 2006; Riès et al., 2013). Data were subsequently re-referenced to an average reference and an independent components analysis (ICA) was performed on each subject in EEGLAB (Delorme and Makeig, 2004). ICA noise components were identified by FASTER criteria (Nolan et al., 2010) as well as spatial correlations of r >0.8 with eyeblink and ballistocardiac artifact templates, and subsequently removed during back-projection.

### 2.6 EEG Periodic and Aperiodic Parameter Estimation

Our analysis pipeline is outlined in Figure 1. Data from all 32 electrode channels were segmented into 2-second epochs corresponding to scan markers for each volume. This yields EEG spectral data and fMRI BOLD data of equal length (182 2-second epochs) for use in a parametric modulation analysis (see below). The power spectrum was obtained within each epoch by computing a Fast Fourier Transform (FFT) with a 2-second Hanning window resulting in a frequency resolution of 0.5 Hz (yielding 98 frequency bins between 1-50Hz). For each resulting power spectra, we applied the methods of FOOOF (Fitting Oscillations and One Over F, https://fooof-tools.github.io/fooof/, Donoghue et al., 2020) within a frequency range of 1-50 Hz to isolate periodic and aperiodic aspects of the spectrum. FOOOF models the power spectrum as a combination of the aperiodic component and a sum of Gaussians (see Figure #2 from Donoghue et al., 2020). As described in Donoghue et al., the aperiodic component is modeled using a Lorentzian function: *L*=*b* - log(*k* + *F*^**β**^), where *F* is a vector of input frequencies, *b* is the spectral offset, **β** is the spectral exponent and *k* is the knee parameter. Fitting was done in ‘fixed’ (*k*=0) rather than ‘knee’ mode, given the relatively narrow range (1-50 Hz) of our frequency data. Adding the ‘knee’ parameter did not change the overall fit (see Supplemental Figure S10). FOOOF utilizes initial seed values for the offset and exponent from the estimate of a first-pass least-squares fit. This fit is subtracted from the original spectrum to generate a flattened spectra (Figure 2B). Subsequently, a power threshold is set (the 2.5th percentile) which identifies the lowest-power points from the residuals that are most likely to reflect the aperiodic part of the spectrum. These points are used in a final estimate of the aperiodic exponent and aperiodic offset values. This fitting procedure performed relatively well in our data: the mean R^2^ value for all subjects, channels and TR intervals was 0.64 (range=0.51-0.74, see Supplemental Figure S1).

**Figure 1.**
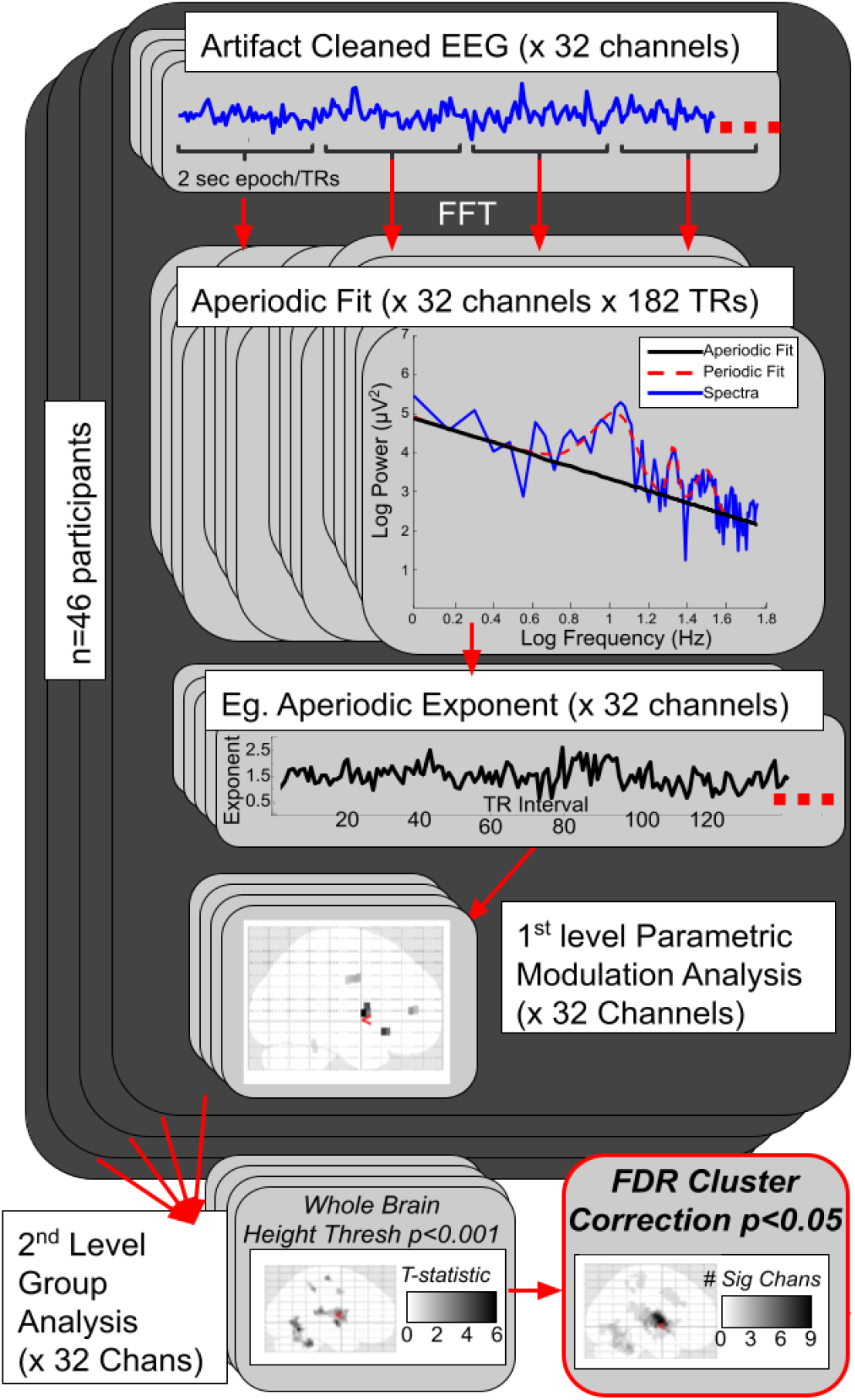
Schematic outline of the analysis pipeline. Artifact Cleaned EEG data (32 channels per participant) from a 6-minute resting-state recording is epoched into 2 second intervals corresponding the fMRI repetition time (TR). A Fast Fourier Transform is performed on each 2 second epoch to yield power spectra (a representative spectra at channel P3 from one participant is shown on the top of the right column). The data is then fit to a power law exponent using FOOOF (see Methods) every 2 seconds to yield a time series for each parameter; spectral exponent, spectral offset, residual alpha power, residual beta power and residual gamma power. After convolution with a standard HRF, the time series is entered as a BOLD predictor in a whole brain parametric modulation analysis (the first level model). A second level model identifies clusters across all subjects that survive a height threshold of p<0.001 and a spatial extent of 10 voxels for each channel (a representative model at channel P3, is shown at with clusters from an SPM glass brain that survive thresholding; see bottom left). Lastly, FDR correction is performed on all clusters from all models/channels (32) and surviving clusters are scored according to the number of times a voxel within a cluster was identified significant on any channel, yielding multi-channel “super-clusters” (see bottom right column).

Residual periodic power was isolated from the spectra by subtracting the final aperiodic fit to create a flattened spectra. Next, we extracted mean values from the flattened spectra within predefined frequency ranges for use in the parametric modulation analysis. We utilized the EEG band designations of a prior simultaneous EEG-fMRI study that included all frequency bands (Mantini et al., 2007) and are consistent with definitions provided by the International Federation of Clinical Neurophysiology (Babiloni et al., 2020). We adjusted our beta EEG range to begin at >16 Hz to avoid artifacts introduced by our fMRI sampling rate of 15 Hz (seen as a peak on our EEG spectra). For the subsequent parametric modulation analysis we only analyzed frequency bands for which the FOOOF algorithm could identify putative periodic peaks for the majority of epochs (see Supplemental Figure S2). On average, periodic peaks were identified in 2.4% of epochs over all participants for delta (1-3 Hz, range of participants: 0.2-7.3), 29.7% for theta (4-7 Hz, range of participants: 10.3-61.1), 70.6% for alpha (8-12 Hz, range of participants: 35.0-93.8), 77.7% for beta (17-25 Hz, range of participants: 35.4-93.9) and 77.1% for low gamma (30-50 Hz, range 55.1-91.8).

### 2.7 EEG-fMRI analysis

Time series corresponding to each two second TR for each EEG parameter of interest (aperiodic exponent, aperiodic offset, residual alpha, residual beta and residual gamma power) were z-scored. Epochs rejected based on FASTER preprocessing were replaced with zeros so as to have no influence on the estimated beta maps. We employed a whole-brain, parametric modulation analysis (Büchel et al., 1998) to estimate BOLD dynamics from the dynamics of each of the EEG parameters under study (Laufs et al., 2003b). Such analyses convolve an assumed HRF with condition specific variables (the parametric modulators) to determine if the BOLD positively or negatively covaries with the modulator variable. In this study, the resulting EEG parameter time series were used as parametric modulators for first-level fMRI analyses in SPM8. aCompCor noise components and motion parameters were also included as nuisance regressors in the general linear model (GLM). For each EEG parameter, beta coefficients were estimated from separate GLM’s for each voxel’s time series, resulting in beta images representing BOLD-EEG parameter associations. Beta images were normalized to the Montreal Neurological Institute’s EPI template (http://www.bic.mni.mcgill.ca) using each subjects’ mean functional image following realignment and smoothed with a 6 mm Gaussian kernel.

Second-level group models identified EEG-fMRI clusters that showed main effects of each parametric modulator. Our initial voxelwise cluster-finding threshold was set to p=0.001 (two-tailed) and a spatial extent of 10 voxels. For each EEG parameter, all clusters surviving the initial whole-brain height threshold, for any of the 32 channels, were retained and these cluster-level p-values were FDR corrected to account for multiple comparisons. A representative single channel model and cluster correction procedure is shown in Supplemental Figure S3. All remaining clusters after FDR correction were saved as binary masks. These 32 masks were summed to create “super-clusters” representing significant clusters of voxels predicted by the EEG parameter for at least one electrode. The numerical weight at each voxel within a supercluster indicates the number of EEG electrodes that identified that voxel within a significant cluster (see also, Figure 1).

## 3. RESULTS

### 3.1 Aperiodic Characteristics of Resting EEG during simultaneous fMRI acquisition

Similar to others, we found that the log-transformed EEG power spectra shows aperiodic or scale-free-like characteristics (Figure 2A). The average spectral offset across all subjects and channels was 5.19±0.36 (log μV^2^, range 4.36-6.29) and the average spectral exponent was 1.49±0.17 (range 1.18-1.91). Both the spectral offset and exponent varied significantly across electrode location (Figure 2C repeated measures ANOVA; offset: F(1,45)=9799.7, p<0.0001; exponent: F(1,45)= 3490.1, p<0.0001; see Supplemental Figure S12 for post-hoc tests). We had no *a-priori* hypothesis regarding the scalp distribution of aperiodic parameters, so retained all channels for subsequent parametric modulation analyses with the fMRI signal. Similar to Wen and Liu (2016b), we found a robust correlation between the spectral offset and exponent, average *r* coefficient of 0.92±0.02 (range: 0.77-0.98) across all subjects, electrodes and TR epochs.

**Figure 2.**
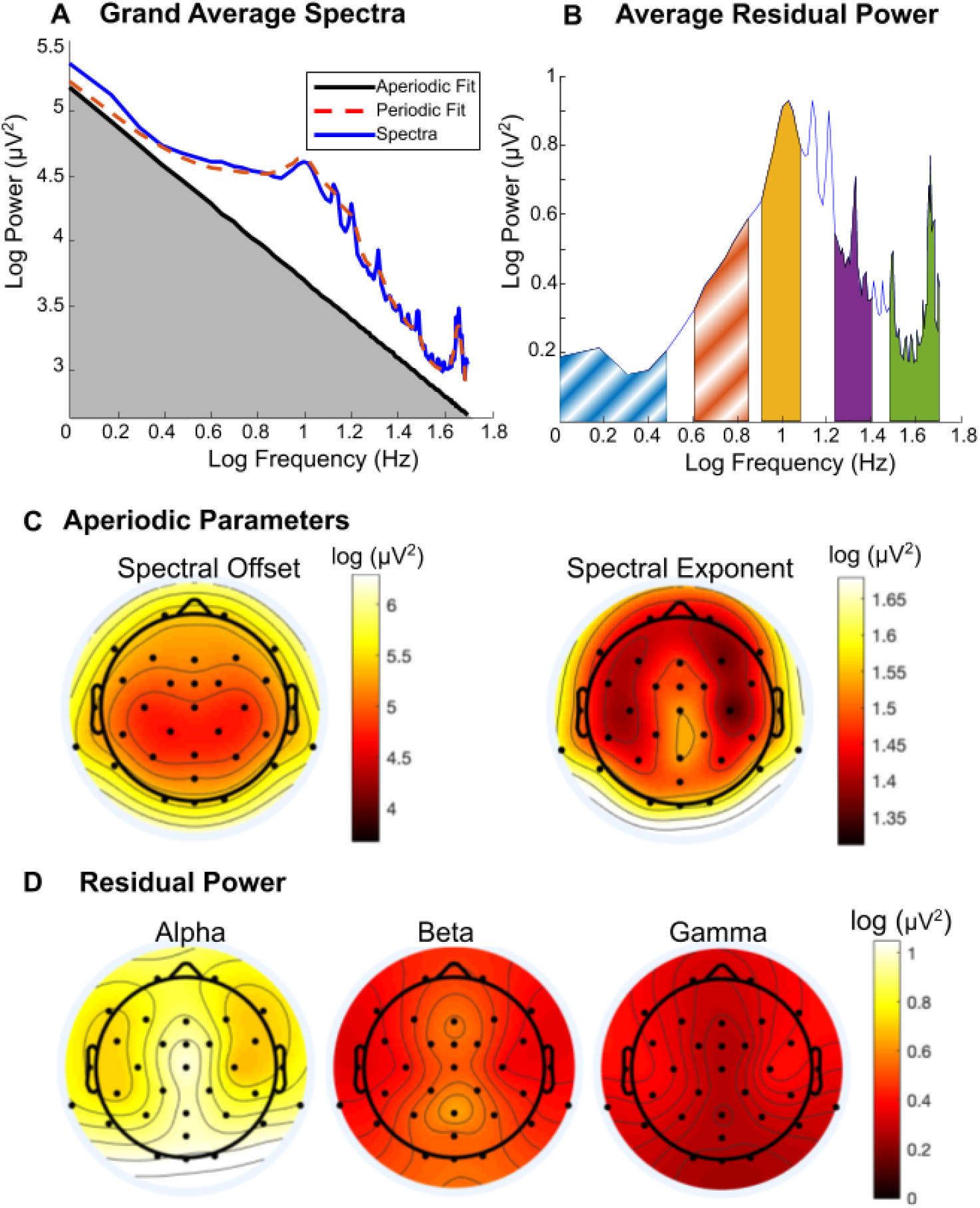
(A) Grand average spectra for all subjects, channels and epochs shown in blue. The average periodic fit output from FOOOF is shown in dashed red and the average aperiodic fit is shown in black. The area in gray under the curve, reflects an estimate of total power related to the scale-free, aperiodic component. (B) Average residual power, after subtracting the aperiodic fit from the average spectra, determined from the average power in the following frequency ranges: delta (0.5-3 Hz, blue), theta (4-7 Hz, red-orange), alpha (8-12 Hz, yellow), beta (17-25 Hz, purple) and gamma (30-50 Hz, green). Delta and theta residual power were not included in subsequent analysis (indicated by white diagonal bars, see Methods). (C) Scalp distribution of aperiodic fit parameters indicates a midline, posterior parietal minimum for the spectral offset power (left) and posterior parietal maximum for the spectral exponent (right). (D) Scalp distribution of residual power reveals a posterior occipital maximum for alpha (left), a central parietal maximum for beta and a bitemporal maximum for gamma.

Average power from the periodic EEG frequency bands was determined after subtracting the aperiodic component to yield ‘flattened’ power spectra (Figure 2B). Residual spectral peaks were consistently detected in the alpha (8-12 Hz), beta (17-25 Hz) and low gamma (30-50 Hz) range (see *Methods*). Average values from the flattened spectra were retained for parametric modulation analyses with the fMRI signal. The alpha band yielded the most residual power (mean±std: 0.85±0.20 log μV^2^, range: 0.44-1.39), followed by beta (mean±std: 0.46±0.13, range: 0.13-0.74) and gamma (mean±std: 0.32± 0.07, range: 0.20-0.46). EEG residual band power revealed a characteristic distribution across the scalp, with alpha power concentrated in posterior/occipital electrodes, beta power concentrated in parietal and central electrodes and low gamma power concentrated bitemporally (Figure 2D).

### 3.2 Parametric Modulation Analysis: Aperiodic Parameters Predict BOLD

#### 3.2.1 Spectral Offset

The aperiodic spectral offset revealed robust correlation with BOLD signals across several brain regions (Figure 3B). Most electrodes (25/32) identified at least one positive cluster corrected for a False Discovery Rate (FDR p<0.05 Figure 3C). Four large (>500 voxels) positive ‘superclusters’ were identified from at least five significant EEG electrodes (see Table 2) and reflect an extended posterior salience network comprised of the bilateral rolandic operculum (including posterior insula and auditory regions, Supplemental Figures S4 and S5), mid-anterior cingulate (Supplemental Figure S6) and the bilateral cerebellum (Supplemental Figure S7). More than half of the electrodes (18/32) identified at least one negative cluster (FDR p<0.05, Figure 3C). One large negative supercluster (~1000 voxels) was identified from 11 electrodes and spanned the superior and medial frontal gyri (Supplemental Figure S8). Several smaller negative clusters (<100 voxels) were widely distributed and reported in Table 2.

**Figure 3.**
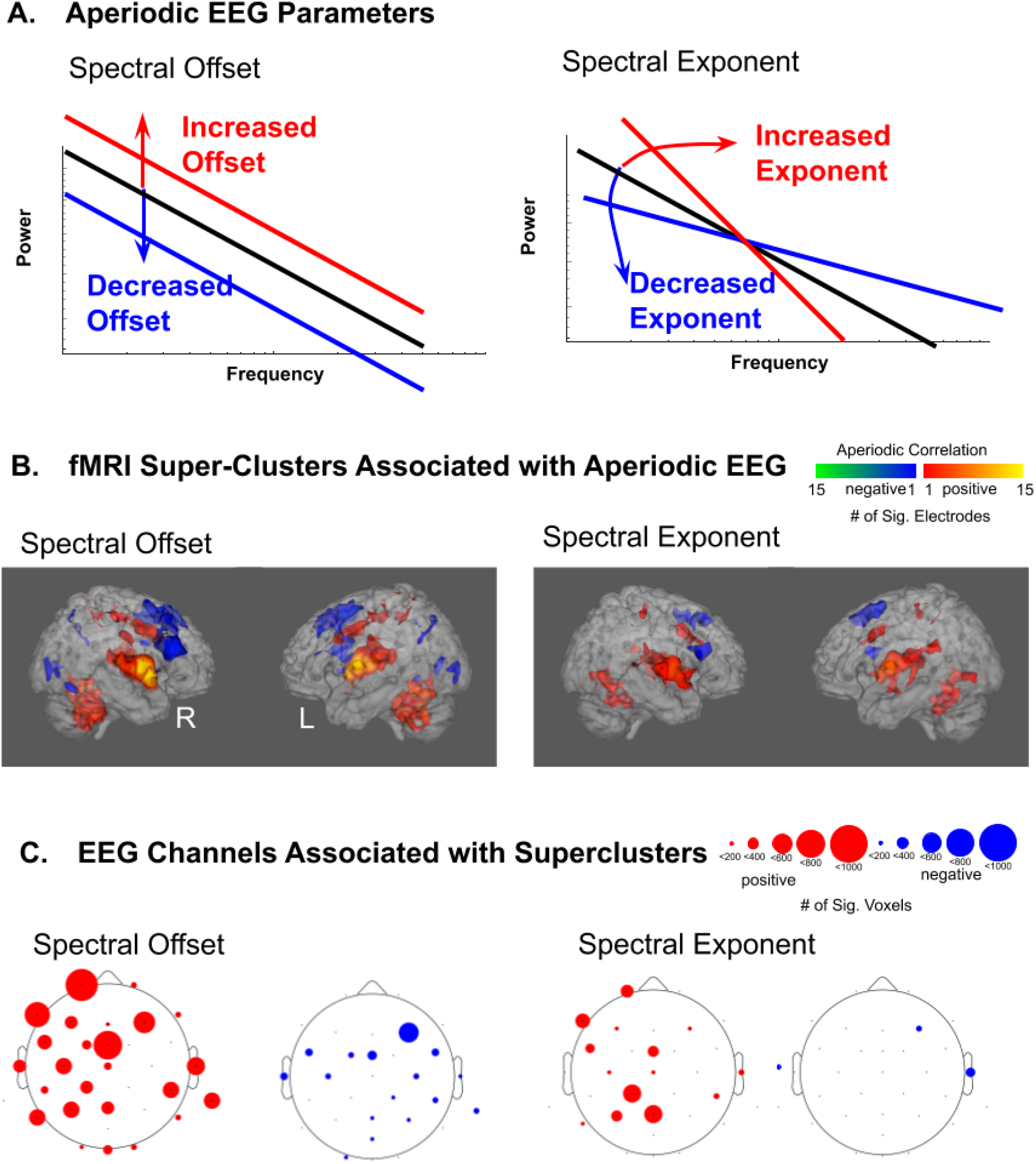
(A) Graphical illustration of the modulation of aperiodic parameters for offset (left) and exponent (right). (B) fMRI Super-Clusters from the multi-channel parametric modulation analysis, identifies regional BOLD activity predicted by EEG the spectral offset (left) and exponent (right). Glass brain illustrations graphically depict the number of EEG electrode channels that predict FDR-corrected fMRI clusters (positively or negatively, as indicated by the color heat map indicated in the legend). (C) EEG Channels that contribute significant FDR-corrected clusters to the multi-channel parametric modulation analysis are shown for offset (left) and exponent (right). The size of the circle at each electrode channel indicates the total number of voxels, after multi-channel FDR-correction, contributed by each channel.

**Table 2.**
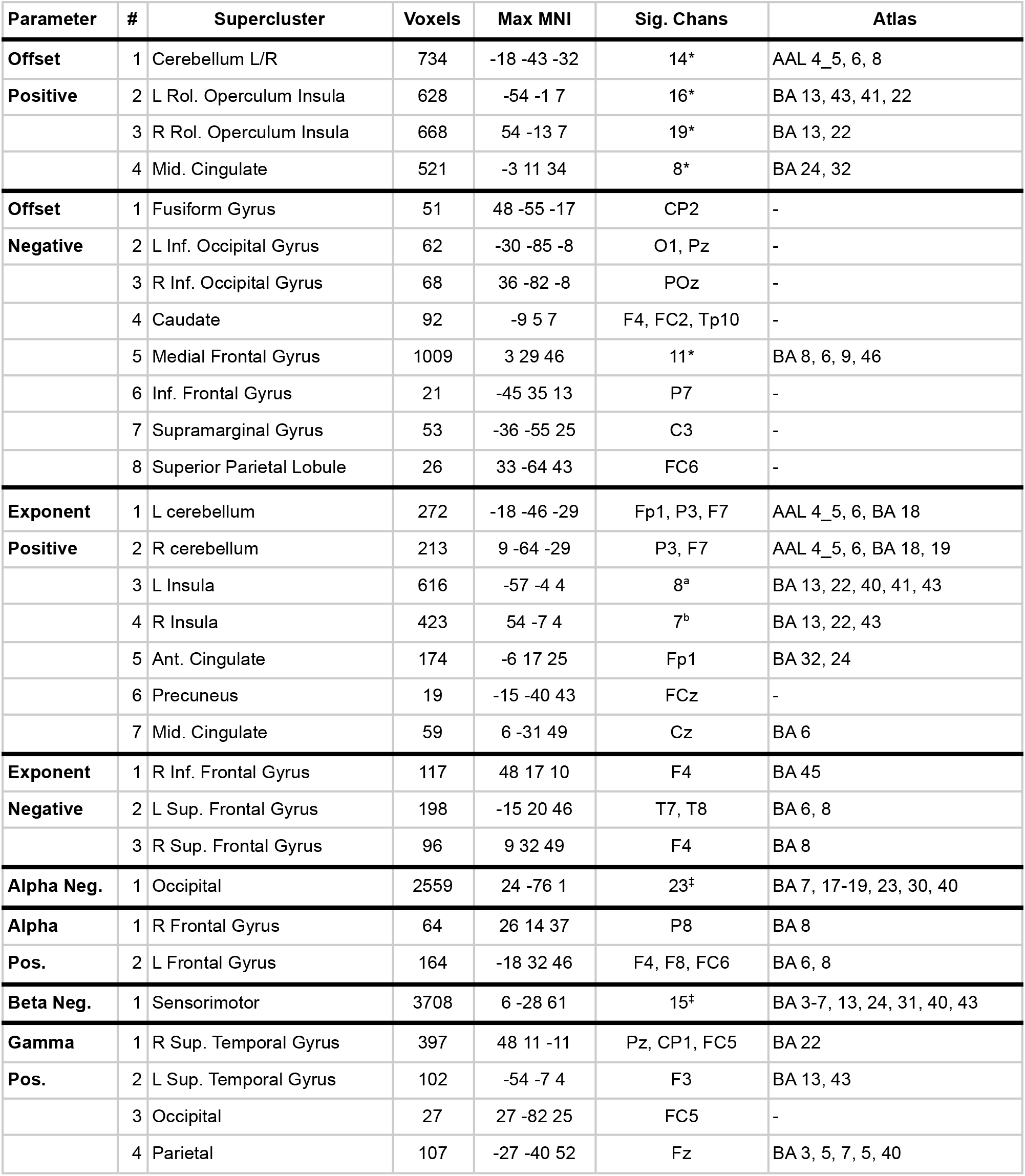
Regions with significant FDR corrected clusters across all 32 EEG channels for each parametric modulator (EEG measure). Brodmann areas are reported for clusters containing more than 20 voxels from that area. *Indicates superclusters with detailed anatomy and EEG channel contributions reported in the Supplemental Figures. ^a.^ F4, C3, P3, T8, Pz, CP1, FC5, CP6; ^b.^Fp1, F3, P7, FCz, Pz, CP1, FC5. ‡Indicates superclusters with EEG channel contributions depicted in Figure 3.

#### 3.2.2 Spectral Exponent

The aperiodic spectral exponent revealed robust correlation with BOLD in many of the same brain regions identified by the spectral offset (Figure 3B), however, fewer significant clusters were retained after correction for multiple comparisons. This was not surprising, given the robust correlation between the exponent and the offset. Slightly less than half of the electrodes (14/32) identified at least one positive cluster (FDR p<0.05, Figure 3C). This analysis reveals five modest superclusters (174-616 voxels) forming an extended network comprising the mid-cingulate, bilateral rolandic operculum (including posterior insula and auditory regions) and the bilateral cerebellum. Only two electrodes identified at least one negative cluster (FDR p< 0.05, Figure 3C), yielding three smaller superclusters (96-198 voxels) including the superior and inferior frontal gyri.

#### 3.2.3 Residual Alpha

Residual Alpha revealed robust negative correlation with BOLD signals in posterior/occipital brain regions (Figure 4B). Most electrodes (24/32) identified at least one negative cluster (FDR p<0.05, Figure 4C), yielding one large negative supercluster (2559 voxels) spanning primarily occipital visual areas, but also some posterior parietal regions (see Table 2). Two smaller positive clusters (64 and 164 voxels) in the bilateral superior frontal gyrus were identified from four EEG electrodes (FDR p<0.05, Figure 4C).

**Figure 4.**
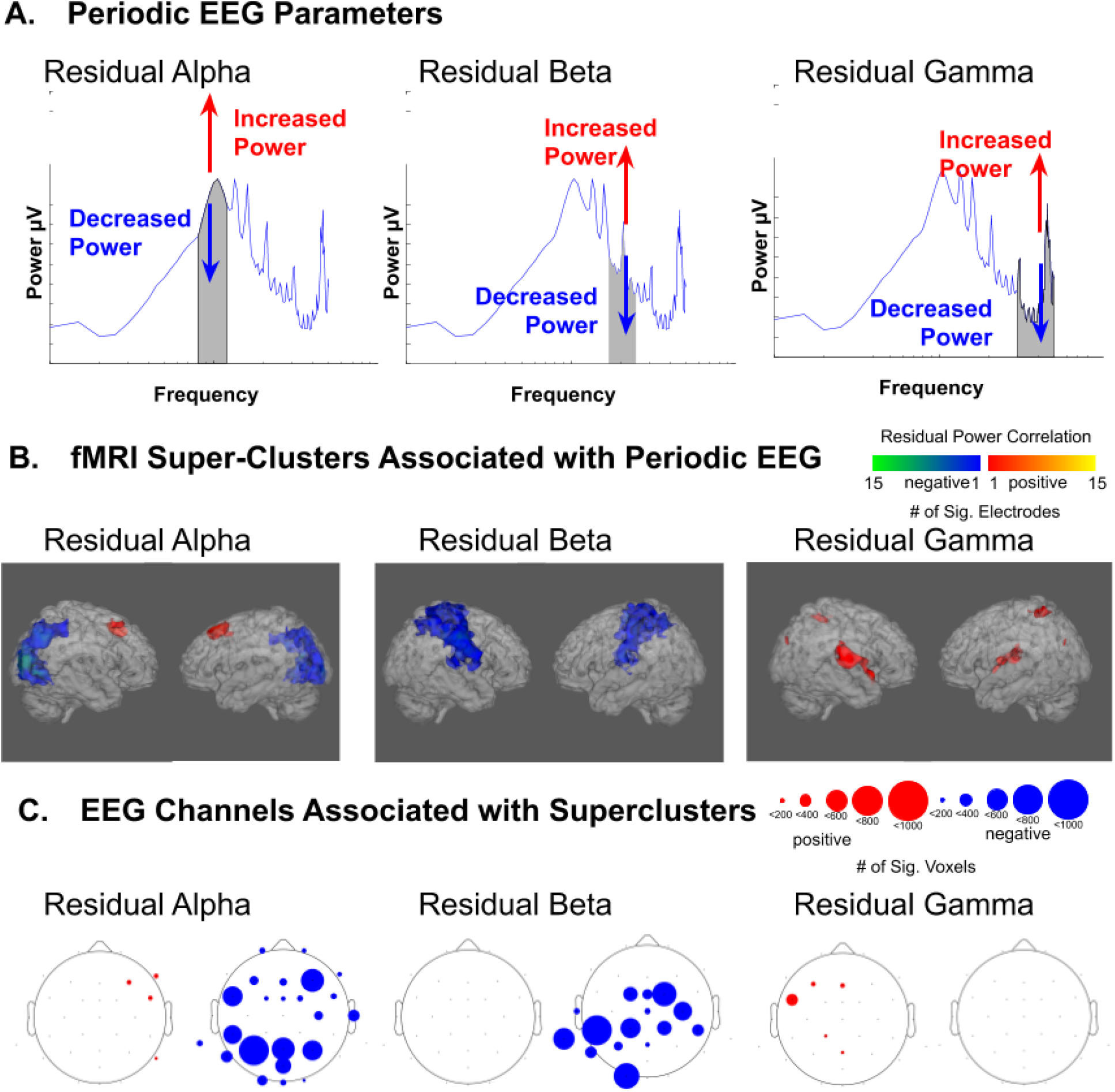
(A) Graphical illustration of the modulation of residual alpha (left), residual beta (center) and residual gamma (right). (B) fMRI Super-Clusters from the multi-channel parametric modulation analysis, identifies regional BOLD activity predicted by EEG residual alpha (left), residual beta (center) and residual gamma (right). Glass brain illustrations graphically depict the number of EEG electrode channels that predict FDR-corrected fMRI clusters (positively or negatively, as indicated by the color heat map indicated in the legend). (C) EEG Channels that contribute significant FDR-corrected clusters to the multi-channel parametric modulation analysis are shown for residual alpha (left), residual beta (center) and residual gamma (right). The size of the circle at each electrode channel indicates the total number of voxels, after multi-channel FDR-correction, contributed by each channel.

#### 3.2.4 Residual Beta

Residual Beta revealed robust negative correlation with BOLD signals in parietal and frontal brain regions around the central gyrus (Figure 4B). Nearly half of the electrodes (15/32) identified at least one negative cluster (FDR p<0.05, Figure 4C), yielding one large negative supercluster (3708 voxels) spanning frontal premotor and motor areas as well as parietal somatosensory areas (see Table 1). No positive clusters were retained after FDR correction across all channels.

#### 3.2.5 Residual Gamma

Residual gamma revealed clusters with positive correlation with BOLD signals primarily in the superior temporal gyrus (figure 4B) as identified from five electrodes (FDR p<0.05, Figure 4C). These clusters were of modest size (102-397 voxels). No negative clusters were retained after FDR correction across all channels.

### 3.3 Aperiodic Parameters Minimally Predict DMN BOLD

Given our hypothesis, based on the work of others (Freeman et al., 2009), that aperiodic EEG would be associated with activity in the DMN, we conducted an ROI analysis of the DMN using a mask from Shirer et al. (2012) We found minimal overlap between the DMN and our EEG-fMRI derived superclusters (see Supplemental Figure S9). 79 voxels (3.1%) of the negative alpha supercluster and 50 voxels (1.3%) of the negative beta supercluster overlapped with DMN in the precuneus. 90 (8.9%) voxels of the aperiodic frontal gyrus negative supercluster overlapped with this literature defined DMN. Thus, a small portion of the anterior hub of the DMN is associated with decreases in aperiodic power and a small portion of the posterior hub of the DMN is associated with decreases in alpha and beta power. These findings provide only minimal evidence to support our initial hypothesis.

## 4. DISCUSSION

In a resting-state, simultaneous EEG-fMRI paradigm, we find that aperiodic EEG variance shows a robust correlation with BOLD hemodynamics. Contrary to our *a-priori* hypothesis, modulation of aperiodic EEG power is only minimally associated with activity in the default mode network (DMN), but instead, predicts activity in anticorrelated, auditory-salience-cerebellar and prefrontal networks. Whereas fMRI-only resting-state studies found anticorrelated signals between frontal-parietal task networks and the DMN (Fox et al., 2005), our main result suggests a prominent role for the salience network, consistent with findings that the salience network coordinates the balance between DMN and executive networks (Goulden et al., 2014). Our findings offer an alternative perspective on resting-brain physiology that considers EEG-BOLD associations: external stimulus processing, not merely “internal” processing within the DMN, drives resting fMRI hemodynamics. We speculate that EEG signals provide a link between the neural response to ongoing environmental stimuli (which are never totally absent in any paradigm) and hemodynamics, which would otherwise be averaged out during standard resting-state, fMRI-only paradigms. We hypothesize that the “environment” in neuroimaging paradigms is the embodied environment in the fMRI, primarily including scanner sound, sensorimotor phenomena, and the explicit instruction not to move. These hypotheses are discussed in detail below.

### 4.1 Comparison with previous literature: EEG-DMN relationships?

Several prior resting-state EEG-fMRI studies have employed the use of the EEG signal as a parametric modulator. Laufs et al. (Laufs et al., 2003b) examined activity in the alpha and beta bands, identifying links to the posterior and anterior cingulate; central hubs of the DMN. This finding appears in direct contrast to our results. We offer several possible explanations for this discrepancy. First, a major difference in our study is the use of eyes open resting state instructions (vs. eyes closed in Laufs et al.). Eye closure has a pronounced effect across EEG bands, increasing alpha and posterior beta power (Barry et al., 2007). In resting-state fMRI, eye closure can increase functional connectivity within the DMN (Weng et al., 2020), whereas resting-state fMRI signals may be more reliable across scan sessions with eyes open instructions (Patriat et al., 2013). We suspect that our use of an eyes open paradigm may have facilitated attention toward the external environment and away from internal “mind wandering,” (Vago and Zeidan, 2016) consistent with greater DMN activity during internal mind-wandering, but not during attention to ongoing stimuli (Van Calster et al., 2017). Moreover, instruction to attend to scanner noise increased activation of auditory and dorsomedial prefrontal cortices in an ROI analysis (Benjamin et al., 2010). Scanner sound may drive arousal (Murphy and Brunberg, 1997), activity in the salience network (Schneider et al., 2016) and scale-free EEG (Lendner et al., 2020); similar to our finding aperiodic EEG activity linked to auditory and salience networks.

Second, we parameterize the EEG spectra to estimate aperiodic and residual periodic components independently, whereas alpha and beta power, as reported in Laufs et al. (Laufs et al., 2003b), likely reflects a combination of periodic and aperiodic activity. Third, we employed a data-driven approach that includes all electrodes, followed by correction for multiple comparisons. Laufs et al. only utilized EEG activity from 2 averaged electrodes (O1 and O2) in their initial model, potentially restricting their findings.

Subsequent work has not deviated much from DMN, ROI-focused analyses and investigations of limited band power, mostly alpha power (Bowman et al., 2017; Goldman et al., 2002; Laufs et al., 2003a; Mayhew and Bagshaw, 2017; Moosmann et al., 2003; Scheeringa et al., 2012). As we have found, resting alpha is highly coupled to BOLD responses in the visual cortex (Feige et al., 2005; Laufs et al., 2006; Mo et al., 2013; Scheeringa et al., 2012), and resting beta is linked to sensorimotor activity (Engel and Fries, 2010; Parkes et al., 2006; Tsuchimoto et al., 2017; Zhang et al., 2008). Using ICA derived fMRI maps, during an eyes-closed paradigm, Mantini et al. (2007) found a mixture of each of the five canonical frequency bands correlated with BOLD in multiple resting-state networks, including a dorsal attention network and auditory cortices. Their finding is broadly consistent with our results: much of the hemodynamic association with resting-state EEG is likely broadband. A key distinction from our results is their identification of the anterior and posterior hubs of the DMN associated predominantly with gamma and beta power, respectively. As noted above, we suspect that these differences are attributable to their use of eyes-closed instructions for the recording period that may alter vigilance and the frequency composition of resting activity (Weng et al., 2020; Wong et al., 2016). We extend these findings to identify links between aperiodic, broadband EEG and ongoing hemodynamic signals.

Few studies have made use of aperiodic or scale-free approaches in simultaneous EEG-fMRI. Portnova et al. (2017) employed a related scale-free measure, Hausdorff fractal dimension, averaged across groups of EEG channels and calculated in a narrow range (2-20 Hz). They found widespread correlations across brain areas including the paracentral lobule, inferior frontal gyrus, superior temporal gyrus and superior occipital gyrus, but the limited bandwidth and channel selection make this finding difficult to interpret in relation to our results. Lei et al. (2015) examined scale-free EEG concurrently with fMRI at rest and during sleep, but they too focused on aperiodic exponent and slope within a narrow range (0.06-3 Hz). Recently developed algorithms enable concurrent estimation of aperiodic and periodic parameters including FOOOF(Donoghue et al., 2020) as we have employed, IRASA (Wen and Liu, 2016b) and curve fitting based on a summation of alpha and the 1/f spectrum (Ouyang et al., 2019). Notably, all of these aforementioned algorithms yield high consistency for parameter estimation (Ouyang et al., 2019). Given robust scale-free EEG-only findings in the literature (Colombo et al., 2019; He et al., 2019; Ma et al., 2018; Muthukumaraswamy and Liley, 2018; Podvalny et al., 2015; Voytek et al., 2015), relative ease of scale-free analysis and dearth of simultaneous scale-free EEG-fMRI studies, we hope our findings might prompt more work in this area.

### 4.2 Interpretation of aperiodic EEG and hemodynamic correlates

Aperiodic and scale-free-like dynamics are frequently attributed to systems of ‘self-organized criticality,’ first described in relation to avalanches in sandpiles (Bak et al., 1987), and thereafter, a myriad of biological and natural phenomena (Bak, 2013). Scale-free activity in the brain has been considered a ‘background’ state, from which linear, rhythmic dynamics emerge to support active processing (Buzsaki, 2006; Freeman et al., 2009). Interpretations of scale-free activity emphasize the presence of long-range-temporal correlations (increase with the aperiodic exponent) which may coordinate activity over multiple spatial and temporal scales (Cocchi et al., 2017; Freeman, 2005), potentially indicative of redundancy and reduced efficiency in the resting state (He, 2014). The aperiodic exponent increases during sleep (Ma et al., 2018), anesthesia (Colombo et al., 2019), and is correlated with pupil size (Pertermann et al., 2019), supporting the perspective that scale-free activity reflects arousal (Lendner et al., 2020). Active stimulus processing reduces the aperiodic exponent (Podvalny et al., 2015; Pozzorini et al., 2013), reflecting an increase in excitatory/inhibitory balance (Gao et al., 2017).

In our eyes-open, resting-state paradigm, we speculate that arousal and attention to environmental signals (in the case of fMRI, primarily auditory) is variable over the course of a 6 minute resting-state scan. Fluctuations in attention might reflect differential engagement of networks associated with top-down (executive, volitional control of attention) and bottom-up (automatic perceptual “capture” of attention), as first proposed by the biased competition theory of attentional control (Buschman and Miller, 2007; Desimone, 1998; Gómez-Laberge et al., 2016) and this theory has been extended to arousal (Mather and Sutherland, 2011). We apply this interpretation as follows: reduced arousal (reduced excitation and increased EEG aperiodic offset/exponent) reflects hemodynamic activity in bottom-up auditory-salience networks, while increased arousal (excitation and decreased EEG aperiodic offset/exponent) reflects top-down executive selection. Prefrontal, executive regions have long been associated with efficiency of information processing (Anderson et al., 2010; Klouda and Cooper, 1990; Sato et al., 2001) and active stimulus processing (Fuster, 2015). Excitation linked to top-down, prefrontal regions may drive arousal (Arnsten et al., 2012) and “execute” correlated signals, as suggested from animal studies (Avery et al., 2014; Merrikhi et al., 2018). Prior literature supports this perspective, including reductions in scale-free correlations during sustained wakefulness (Meisel et al., 2017) and increased scale-free correlations associated with poor attentional performance (Irrmischer et al., 2018).

Associations between aperiodic EEG activity and bottom-up sensory regions may reflect the 1/f statistics of natural stimuli (Lin et al., 2016) and others have found associations between EEG measures of criticality and activity in the auditory cortex (Fagerholm et al., 2015). However, our paradigm likely induced an overall state of low participant arousal (Tagliazucchi and Laufs, 2014) where ongoing scanner sound wasn’t task relevant. Therefore, increased aperiodic activity (increased inhibition) linked to fMRI signal in the auditory cortex likely reflects active suppression of irrelevant scanner sound (Ahveninen et al., 2017), and/or inhibitory current within the auditory cortex from reduced arousal (Lin et al., 2019). Active inhibition might also explain the aperiodic EEG correlation with regions of the salience network, including the posterior insula and cingulate, a network typically associated with increased arousal (Schneider et al., 2016), and anxiety (Seeley et al., 2007). States of low or high arousal (but not moderate arousal) yield anticorrelated activity between the salience network and prefrontal executive networks (Young et al., 2017), moreover, inhibition of the salience network might prevent bottom-up intrusions during periods of rest or sleep (Ma et al., 2020). Further studies are necessary to confirm that resting-state auditory-salience activity reflects an inhibitory process.

The correlation between cerebellar metabolism and aperiodic EEG was unexpected. Nonetheless, the cerebellum has been implicated in the control of cortical efficiency (Caligiore et al., 2017) and signal coordination across multiple time-scales (Guell et al., 2018), the defining feature of scale-free activity. Notably, cerebellar disruption increases cortical noise (Deverett et al., 2019) and cerebellar stimulation increases EEG complexity, based on multi-scale entropy (Farzan et al., 2016), a measure related to scale-free activity. Beyond motor control, the cerebellum modulates sensory processing (Gao et al., 1996), including the timing of auditory events (Grube et al., 2010), possibly explaining our findings that the auditory-salience network is co-modulated with the cerebellum. Lastly, the cerebellum may function more generally in sleep and arousal (Canto et al., 2017), or general postural adjustments (Cebolla et al., 2016; Drew et al., 2019) that might occur during a resting-state scan.

Unlike a sandpile, the brain is concerned with the ongoing maintenance of dynamic activity to support information signaling *in addition to* the ongoing maintenance of its energy supply. Modeling studies have proposed that the scale-free relationships between EEG frequency bands emerge from energetic constraint (Burroni et al., 2017). Our findings further support this perspective: global neural excitability/arousal (defined from aperiodic EEG), co-fluctuating with top-down and bottom-up networks (defined from fMRI activity), suggests an optimization scheme for interdependent electrical and vascular dynamics (Deacon, 2011). Yoking metabolic and electrical balance to global arousal and signal processing, might be akin to “meta-criticality”(De Domenico et al., 2016; Tognoli and Kelso, 2014) where increases in global arousal and processing efficiency are constrained by the energy supply necessary to support “top-down” attention, excitation and stimulus processing in the first place. Reciprocally, arousal must be balanced by periods of rest. The availability of energetic resources in sensory-salience regions might constrain the inhibition of “bottom-up” signals that would otherwise interrupt restorative rest.

### 4.3 Limitations and future directions

Strikingly, we find little association between the DMN and resting EEG signatures, despite the strength of the DMN signal in single modality PET and fMRI studies. One clear limitation of our approach is that the surface EEG has poor anatomical resolution and inability to pick up activity from specific structures in the depth of the brain; some have argued that intracerebral EEG is necessary to examine the DMN (Jerbi et al., 2010). Electrical dipoles originating from midline structures such as the DMN, may be superimposed or buried in the lateralized EEG activity. Another limitation is that the assumptions of the canonical hemodynamic response function may not apply to aperiodic EEG (He, 2011), and non-standard HRFs may need to be investigated (Feige et al., 2017). The interpretation of our findings must be tempered by the fact that we did not probe the psychological state of our participants, during or after scanning, so we cannot ascertain the extent to which their mentation was guided by internal (thoughts, emotions) and/or external stimuli (sights, sounds, sensations). Future studies might make use of experience sampling, or specific tasks to assay attentional orientation while at rest Despite our attempt to apply best practices for cleaning EEG data collected in the MR environment, certain sources of noise (e.g., head motion) are not correctable without more advanced approaches to both data collection and offline de-noising (e.g., copper loop dummy recordings). Lastly, future studies might consider the use of aperiodic EEG-fMRI in clinical populations where deviations from scale-free are hypothesized to reflect pathological processes (Hesse and Gross, 2014). Reductions in scale free EEG power are seen in aging (Voytek et al., 2015) and preliminarily in schizophrenia (Peterson et al., 2018), and both are conditions associated with alterations in frontal metabolism (Weinberger 1988; Davis et al. 2007) and changes in information processing efficiency. Our findings underscore the advantages to combining EEG and fMRI modalities, not merely for the differences in temporal and spatial resolution, but for the importance of reconciling hemodynamic and electrical perspectives on brain activity.

## Acknowledgements and disclosures

This work was supported by grants from National Institute of Mental Health (R01MH058262-17 to JMF) and the VA (I01CX000497-06 to JMF, IK1CX002089 to MSJ). DHM consults for Boehringer Ingelheim. We gratefully acknowledge the contributions of Charles Duffy, Terrence Deacon, Mani Hamidi, Elhum Shamshiri and Parham Pourdavood, who provided helpful critiques on an earlier version of this manuscript. The authors have declared that there are no conflicts of interest in relation to the subject of this study.

