## Supplemental Figure for "Aperiodic measures of neural excitability are associated with anticorrelated hemodynamic networks at rest: a combined EEG-fMRI study"

### SUPPLEMENTAL FIGURES

#### FOOOF Fitting $R^2$ Values

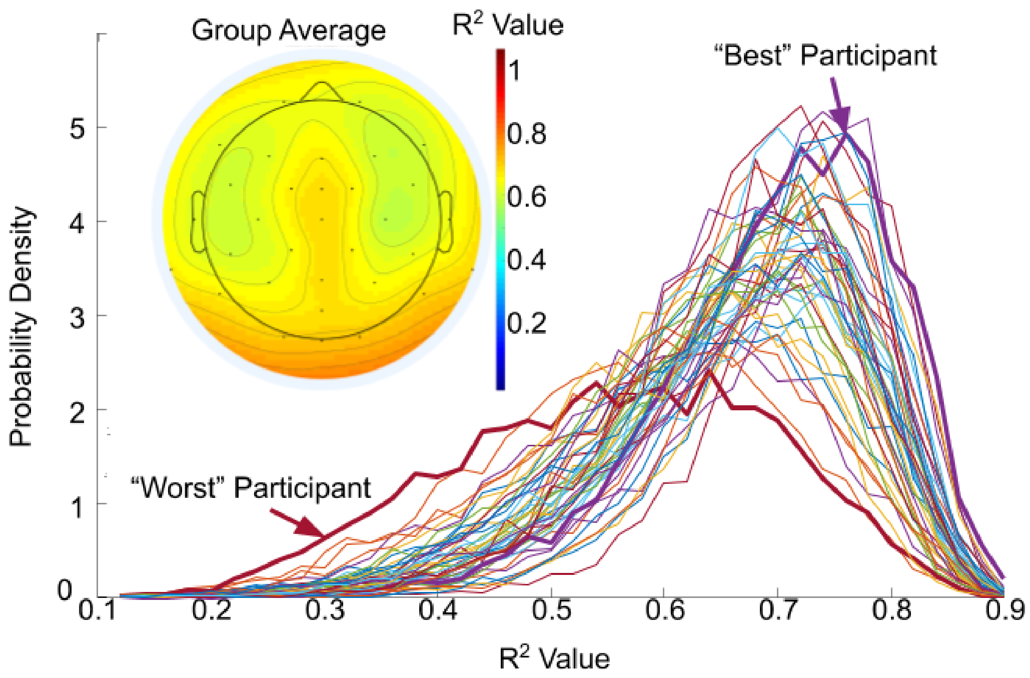


**Supplemental Figure 1.** The distribution of fits for each subject as a probability density function across all channels and TRs (5,824 "fits" per participant) reveals that all participants and TR epochs were fairly well fit by the FOOOF algorithm: mean $R^2$ is 0.64 (mean range for subjects=0.51-0.74) with no clear outlier participants. *Inset*: Distribution of $R^2$ values across the scalp and averaged over all participants also indicates that all electrodes were fit fairly well by the FOOOF algorithm.

**FOOOF Identification of Oscillatory Peaks**

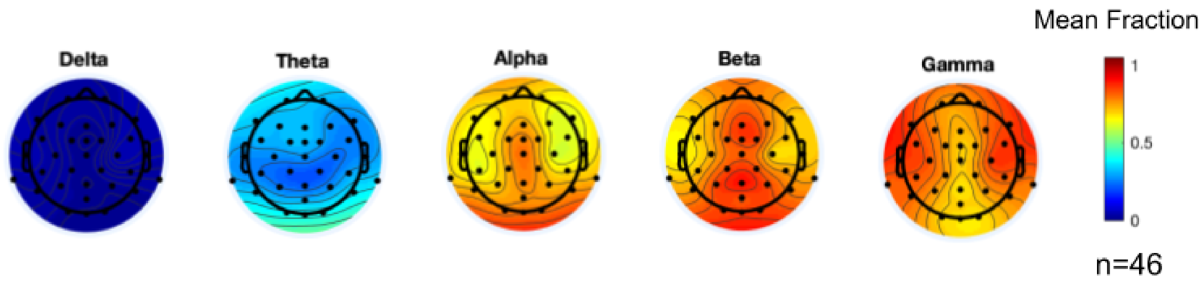


**Supplemental Figure 2.** The fraction of oscillatory peaks identified by the FOOOF algorithm, on average, for all participants and plotted across the scalp for each electrode.

**Example: Single Channel Clusters vs. Multichannel Superclusters**

| # | Cluster Corrected (single channel) | FDR correction (32 channels) |
|---|---|---|
| 1 | 1.02E-14 | 9.40E-13 |
| 2 | 0.02829341 | 0.11082741 |
| 3 | 0.05237058 | 0.18531129 |
| 4 | 0.61110691 | 0.74506179 |
| 5 | 0.3228336 | 0.54641185 |
| 6 | 0.01951591 | 0.08549827 |
| 7 | 0.07804166 | 0.25642259 |
| 8 | 0.65597832 | 0.74506179 |
| 9 | 0.4491182 | 0.6773586 |
| 10 | 0.48644635 | 0.69926663 |
| 11 | 0.21154493 | 0.46505256 |
| 12 | 0.70171995 | 0.74766539 |
| 13 | 0.41410312 | 0.64307725 |
| 14 | 0.65597832 | 0.74506179 |
| 15 | 0.65597832 | 0.74506179 |

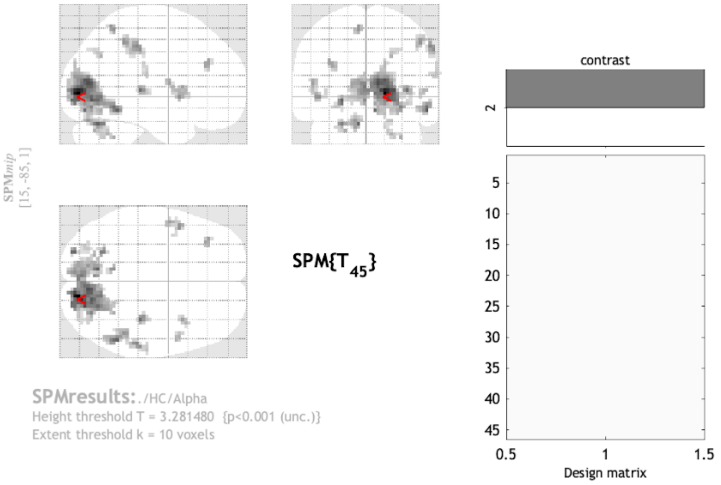


**Statistics:** ***p-values adjusted for search volume***

| set-level p | set-level c | cluster-level $p_{FWE-corr}$ | cluster-level $q_{FDR-corr}$ | cluster-level $k_E$ | cluster-level $p_{uncorr}$ | peak-level $p_{FWE-corr}$ | peak-level $q_{FDR-corr}$ | peak-level T | peak-level $(Z_{\equiv})$ | peak-level $p_{uncorr}$ | mm mm mm |
|---|---|---|---|---|---|---|---|---|---|---|---|
| 0.000 | 15 | 0.000 | 0.000 | 732 | 0.000 | 0.002 | 0.027 | 6.29 | 5.30 | 0.000 | 15 -85 1 |
| | | | | | | 0.085 | 0.200 | 5.16 | 4.55 | 0.000 | 9 -76 10 |
| | | | | | | 0.085 | 0.200 | 5.16 | 4.55 | 0.000 | 24 -79 4 |
| | | 0.028 | 0.037 | 51 | 0.003 | 0.168 | 0.200 | 4.90 | 4.36 | 0.000 | -9 -67 -26 |
| | | | | | | 0.381 | 0.240 | 4.54 | 4.10 | 0.000 | -24 -64 -26 |
| | | | | | | 0.685 | 0.374 | 4.21 | 3.84 | 0.000 | -12 -76 -23 |
| | | 0.052 | 0.052 | 43 | 0.005 | 0.172 | 0.200 | 4.89 | 4.35 | 0.000 | 54 -49 -14 |
| | | 0.611 | 0.293 | 13 | 0.095 | 0.202 | 0.200 | 4.82 | 4.31 | 0.000 | -24 -88 -5 |
| | | 0.323 | 0.214 | 21 | 0.039 | 0.260 | 0.200 | 4.72 | 4.23 | 0.000 | 36 -28 4 |
| | | 0.020 | 0.037 | 56 | 0.002 | 0.485 | 0.282 | 4.42 | 4.01 | 0.000 | 57 -31 43 |
| | | | | | | 0.647 | 0.363 | 4.25 | 3.87 | 0.000 | 48 -43 46 |
| | | 0.078 | 0.062 | 38 | 0.008 | 0.607 | 0.344 | 4.29 | 3.91 | 0.000 | -45 8 -2 |
| | | | | | | 0.772 | 0.446 | 4.11 | 3.77 | 0.000 | -51 -1 10 |
| | | | | | | 0.999 | 0.824 | 3.46 | 3.24 | 0.001 | -39 2 4 |
| | | 0.656 | 0.293 | 12 | 0.108 | 0.656 | 0.363 | 4.24 | 3.87 | 0.000 | -9 -52 58 |
| | | 0.449 | 0.255 | 17 | 0.060 | 0.782 | 0.446 | 4.10 | 3.76 | 0.000 | -36 35 28 |
| | | 0.486 | 0.256 | 16 | 0.067 | 0.795 | 0.446 | 4.08 | 3.74 | 0.000 | 36 11 7 |
| | | 0.212 | 0.152 | 26 | 0.024 | 0.901 | 0.516 | 3.93 | 3.62 | 0.000 | 30 -52 -38 |
| | | | | | | 0.927 | 0.529 | 3.88 | 3.58 | 0.000 | 21 -58 -35 |
| | | | | | | 0.984 | 0.688 | 3.68 | 3.42 | 0.000 | 21 -70 -23 |
| | | 0.702 | 0.310 | 11 | 0.122 | 0.915 | 0.516 | 3.90 | 3.60 | 0.000 | 51 8 4 |
| | | 0.414 | 0.255 | 18 | 0.054 | 0.917 | 0.516 | 3.90 | 3.60 | 0.000 | 21 -64 34 |
| | | | | | | 0.999 | 0.824 | 3.46 | 3.24 | 0.001 | 27 -70 40 |
| | | 0.656 | 0.293 | 12 | 0.108 | 0.987 | 0.702 | 3.66 | 3.40 | 0.000 | 63 -22 19 |
| | | 0.656 | 0.293 | 12 | 0.108 | 0.988 | 0.702 | 3.65 | 3.40 | 0.000 | -30 -79 -11 |

*table shows 3 local maxima more than 8.0mm apart*

Height threshold: T = 3.28, p = 0.001 (1.000)
Extent threshold: k = 10 voxels, p = 0.139 (0.748)
Expected voxels per cluster, <k> = 4.801
Expected number of clusters, <c> = 1.38
FWEp: 5.351, FDRp: 6.293, FWEc: 51, FDRc: 51

Degrees of freedom = [1.0, 45.0]
FWHM = 10.6 10.7 11.4 mm mm mm; 3.5 3.6 3.8 {voxels}
Volume: 1040472 = 38536 voxels = 601.7 resels
Voxel size: 3.0 3.0 3.0 mm mm mm; (resel = 47.70 voxels)

**Supplemental Figure S3.** SPM group (2nd level) results for a single channel (Pz) in the model where residual alpha EEG power is input as a parametric modulator. *Table:* Cluster corrected P-values for the single model are shown alongside 32 channel corrected P-values that were retained for the supercluster analysis reported in the main manuscript.

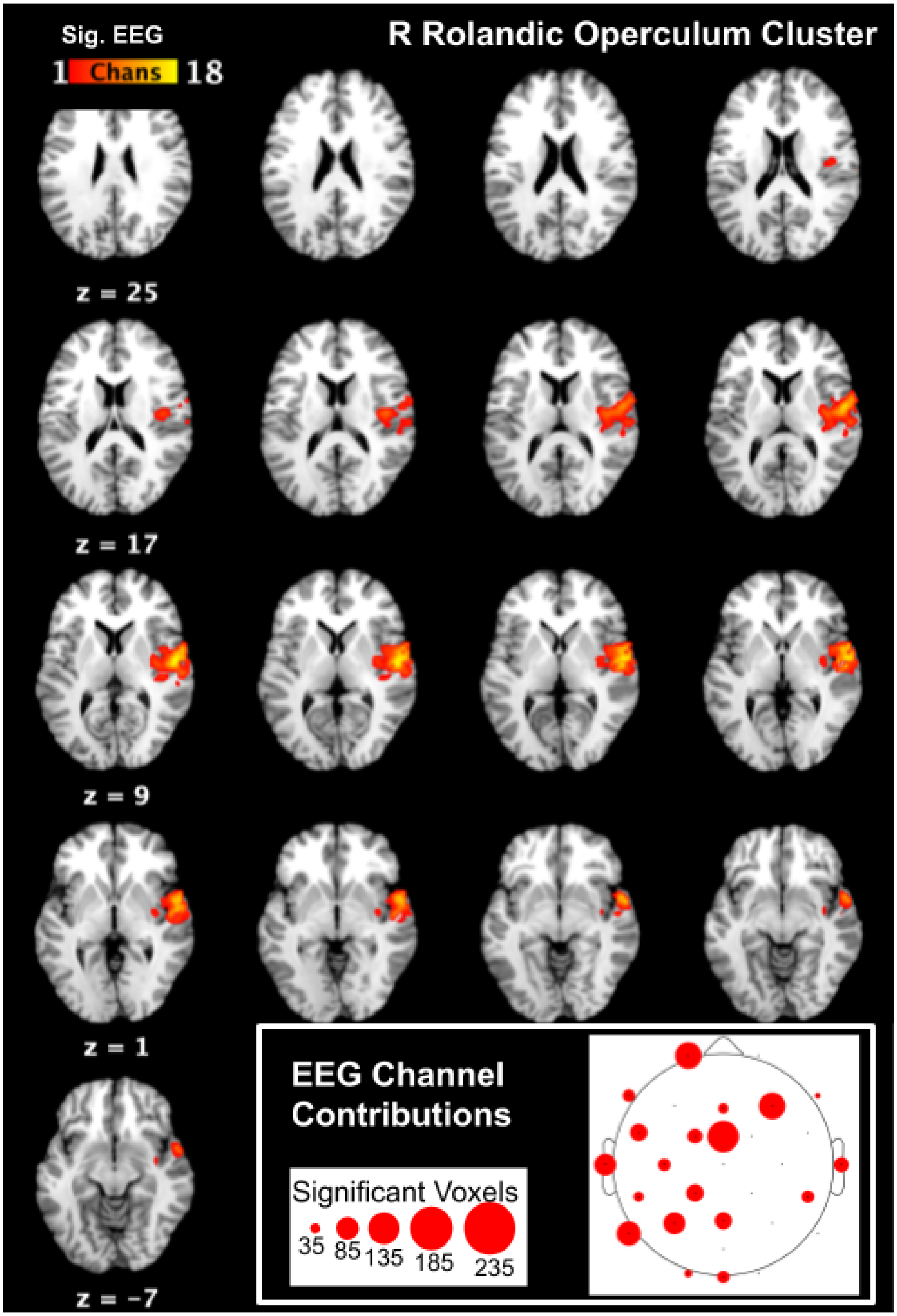


**Supplemental Figure S4.** Axial slices and anatomical distribution of significant voxels that form the R rolandic opercular supercluster. *Inset*: Contribution of significant voxels from EEG channels. The size of the plotted circle on the EEG channel scalp map indicates the approximate number of significant voxels identified at that channel to contribute to the supercluster.

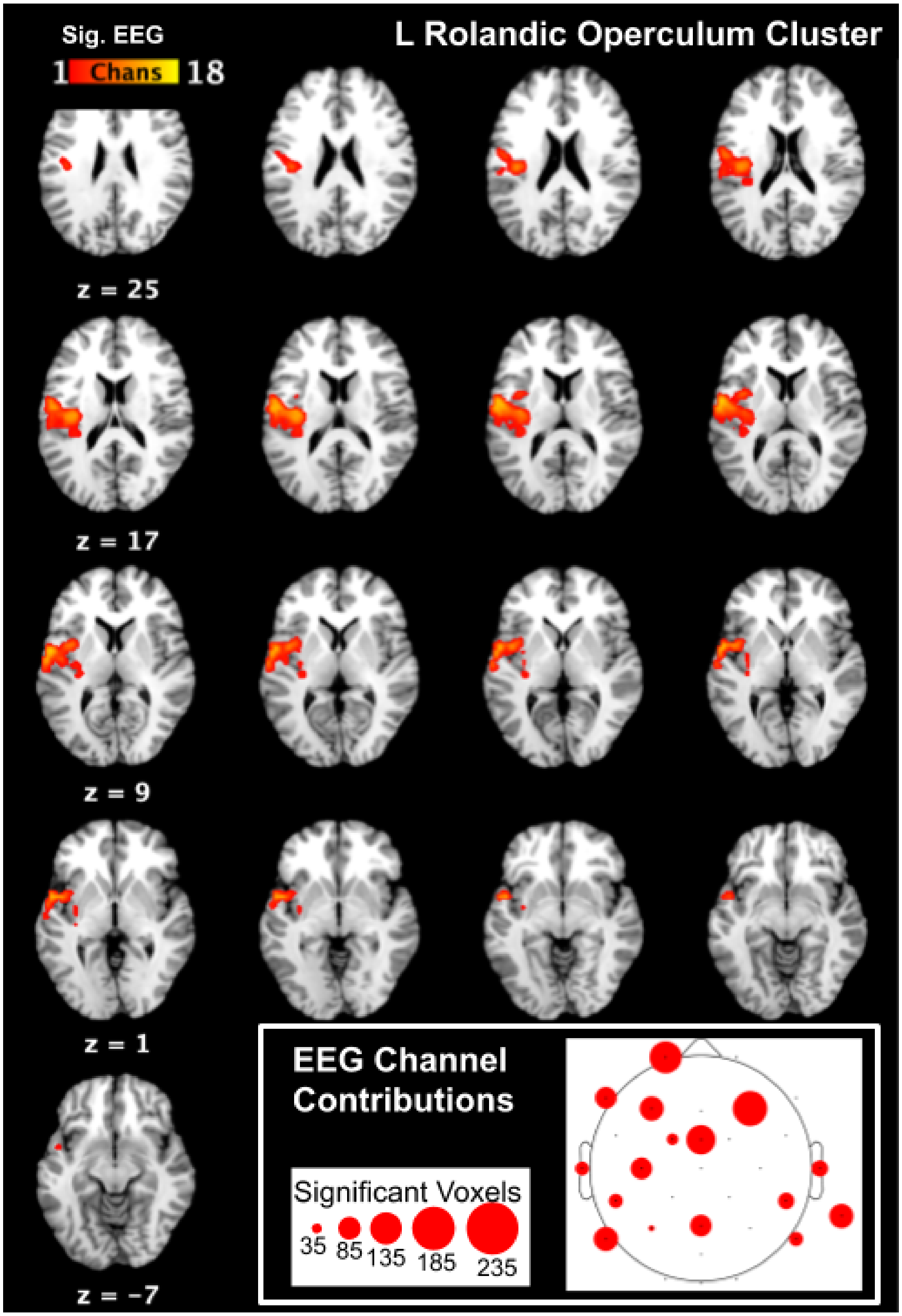


**Supplemental Figure S5.** Axial slices and anatomical distribution of significant voxels that form the L rolandic opercular supercluster. *Inset*: Contribution of significant voxels from EEG channels. The size of the plotted circle on the EEG channel scalp map indicates the approximate number of significant voxels identified at that channel to contribute to the supercluster.

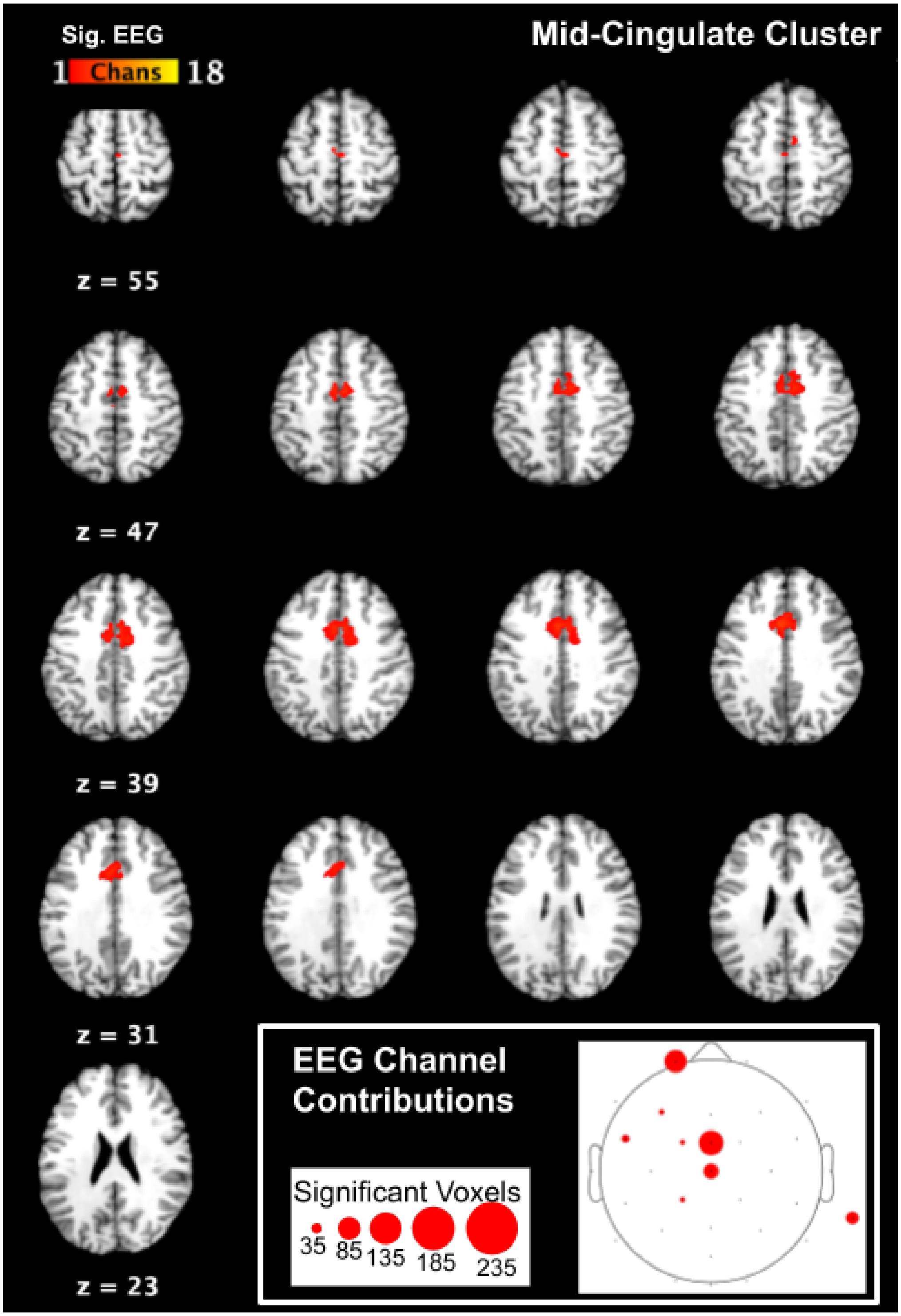


**Supplemental Figure S6.** Axial slices and anatomical distribution of significant voxels that form the mid-cingulate supercluster. *Inset*: Contribution of significant voxels from EEG channels. The size of the plotted circle on the EEG channel scalp map indicates the approximate number of significant voxels identified at that channel to contribute to the supercluster.

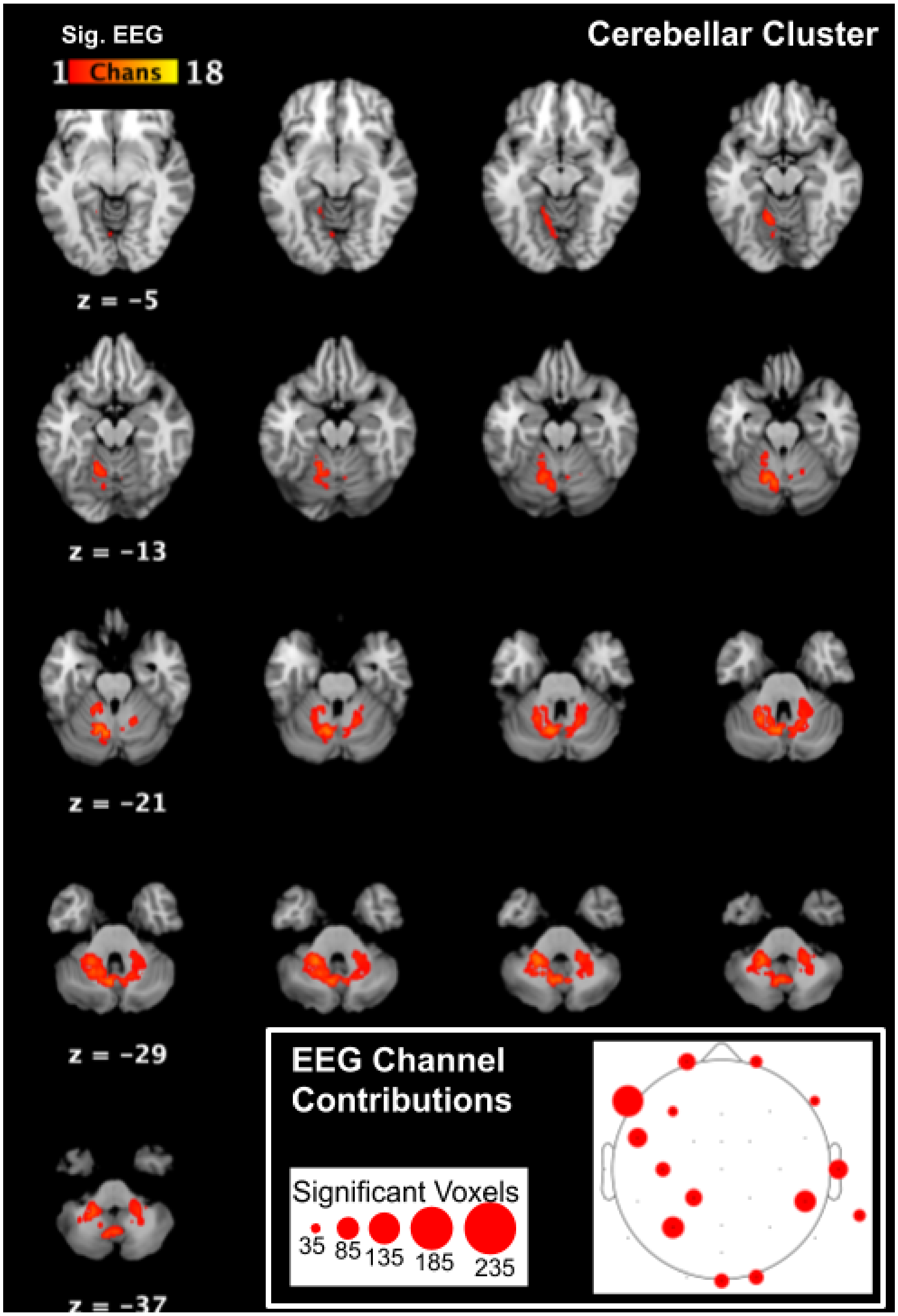


**Supplemental Figure S7.** Axial slices and anatomical distribution of significant voxels that form the cerebellar supercluster. *Inset*: Contribution of significant voxels from EEG channels. The size of the plotted circle on the EEG channel scalp map indicates the approximate number of significant voxels identified at that channel to contribute to the supercluster.

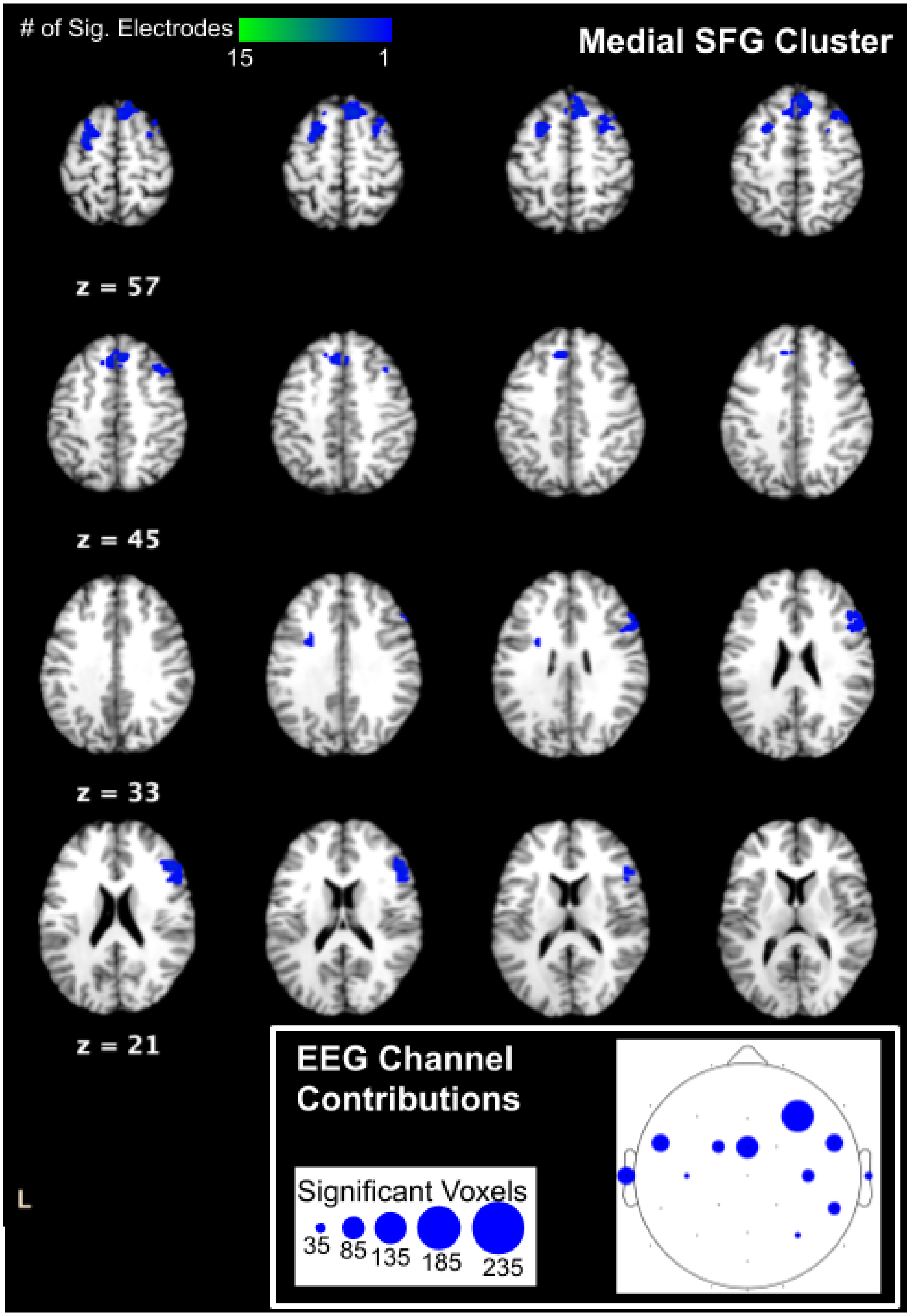


**Supplemental Figure S8.** Axial slices and anatomical distribution of significant voxels that form the medial SFG supercluster. *Inset*: Contribution of significant voxels from EEG channels. The size of the plotted circle on the EEG channel scalp map indicates the approximate number of significant voxels identified at that channel to contribute to the supercluster.

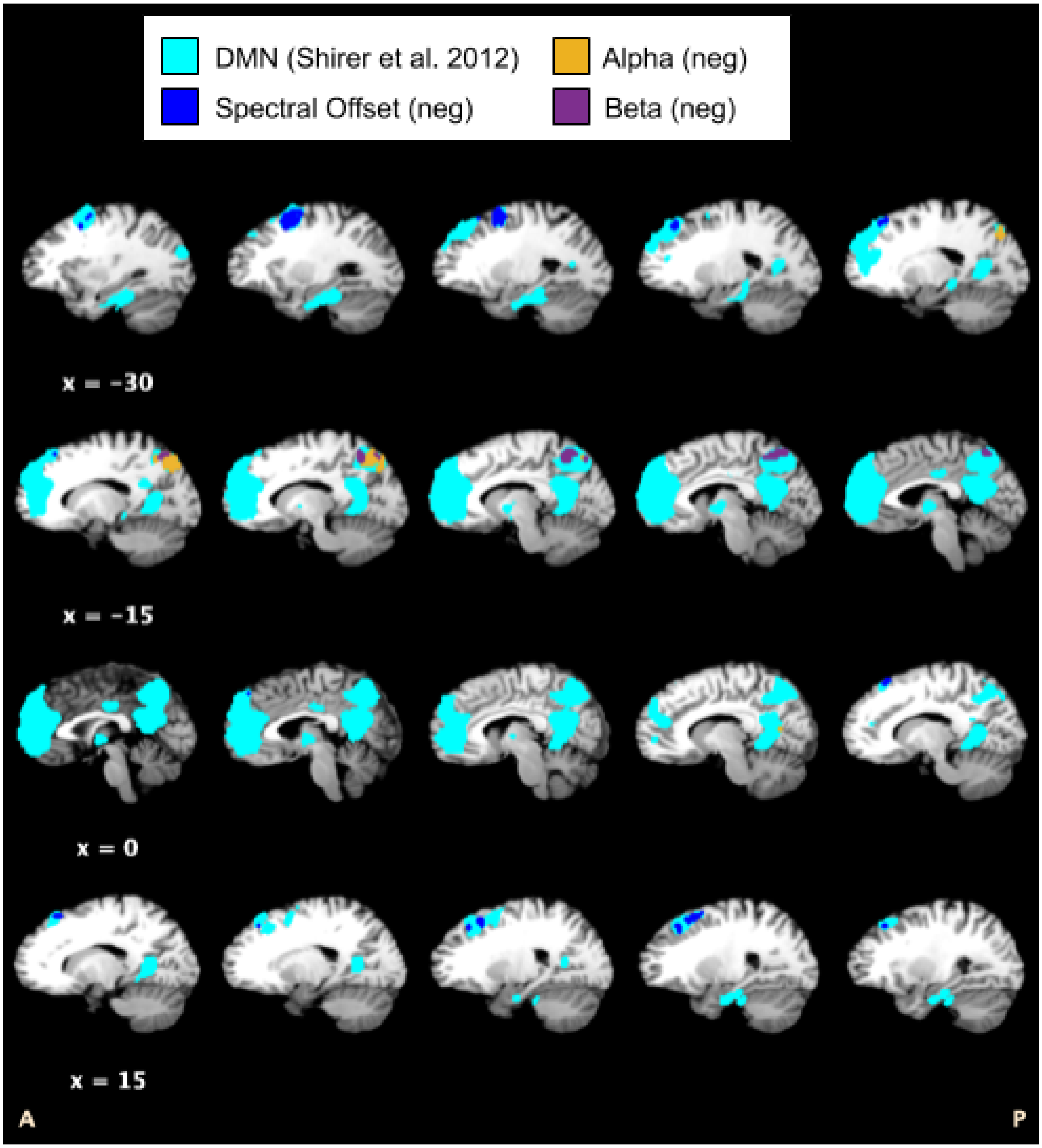


**Supplemental Figure S9.** Supercluster regions (spectral offset, dark blue; alpha, orange; beta, purple) that overlap with a literature defined DMN (cyan), includes dorsal and ventral DMN networks from Shirer et al. 2012.

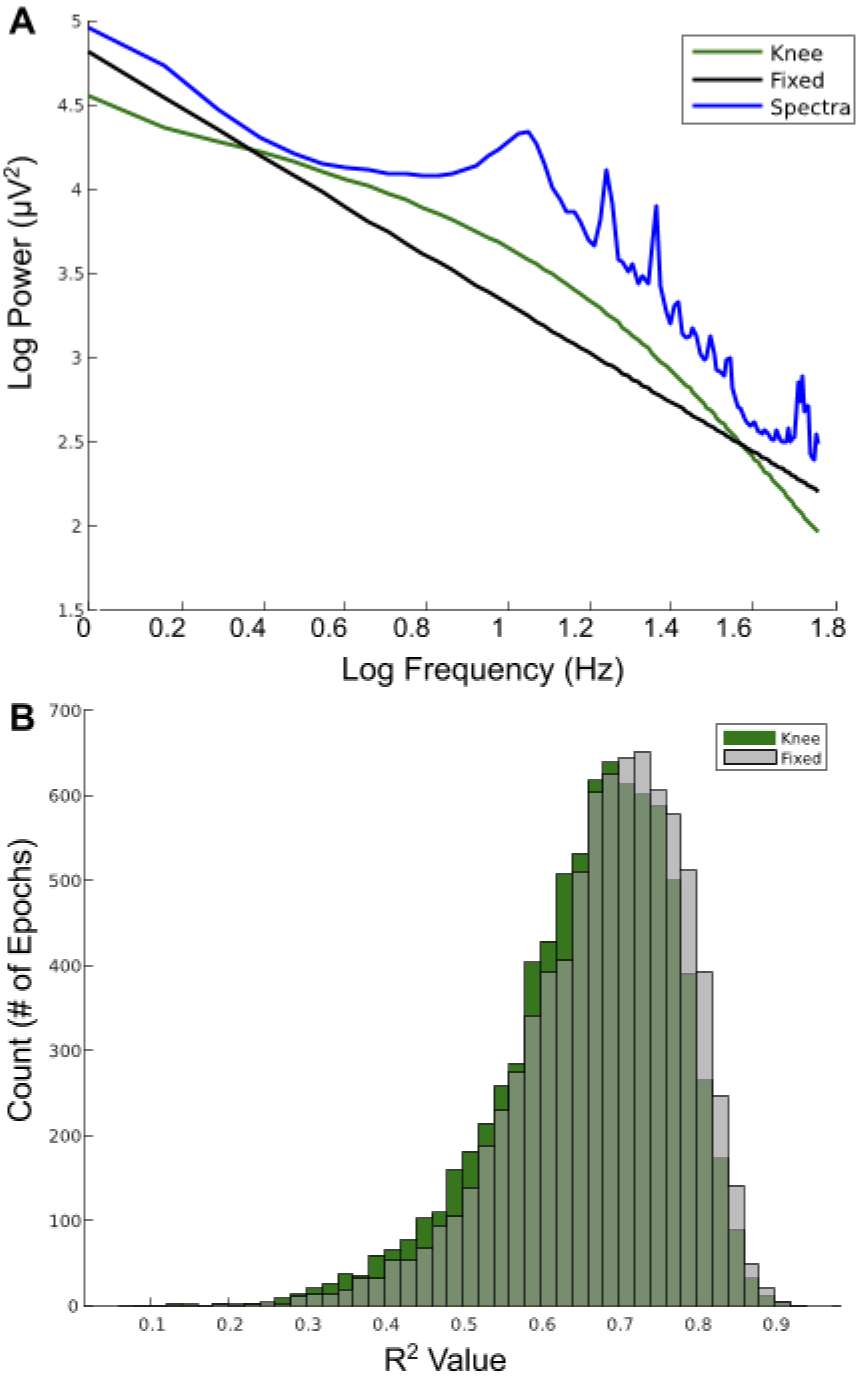


**Supplemental Figure S10. A.** Grand average spectra for an exemplary channel (Pz) for all subjects and epochs shown in blue. The average aperiodic fit output from FOOOF in 'fixed' mode is shown in black. The average aperiodic fit output from FOOOF in 'knee' mode is shown in green. **B.** The distribution of $R^2$ values for FOOOF fits within each 2 second epoch for all subject in 'fixed' mode (gray) and 'knee' mode (in green).

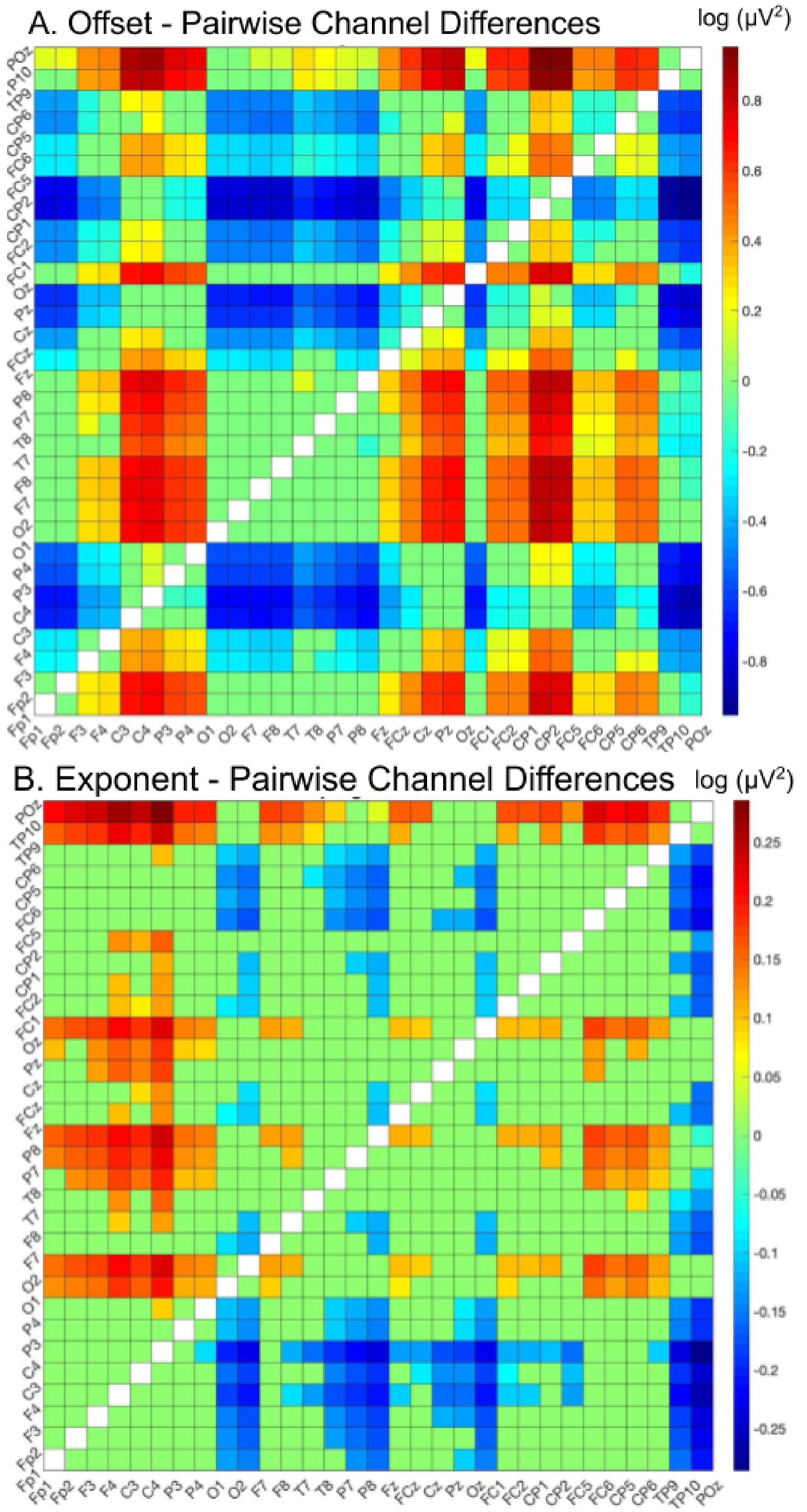


**Supplemental Figure S11.** Significant pairwise differences between all EEG channels. Differences for which the results of post-hoc, multiple comparisons testing (Tukey-Kramer correction) yield a p-value > 0.05 are set to 0. **(A)** Aperiodic Offset **(B)** Aperiodic Exponent.
